# Behavioral and neural representations of spatial directions across words, schemas, and images

**DOI:** 10.1101/220137

**Authors:** Steven M. Weisberg, Steven A. Marchette, Anjan Chatterjee

## Abstract

Modern spatial navigation requires fluency with multiple representational formats, including visual scenes, signs, and words. These formats convey different information. Visual scenes are rich and specific, but contain extraneous details. Arrows, as an example of signs, are schematic representations in which the extraneous details are eliminated, but analog spatial properties are preserved. Words eliminate all spatial information and convey spatial directions in a purely abstract form. How does the human brain compute spatial directions within and across these formats? To investigate this question, we conducted two experiments on men and women: a behavioral study that was preregistered, and a neuroimaging study using multivoxel pattern analysis of fMRI data to uncover similarities and differences among representational formats. Participants in the behavioral study viewed spatial directions presented as images, schemas, or words (e.g., “left”), and responded to each trial, indicating whether the spatial direction was the same or different as the one viewed previously. They responded more quickly to schemas and words than images, despite the visual complexity of stimuli being matched. Participants in the fMRI study performed the same task, but responded only to occasional catch trials. Spatial directions in images were decodable in the intraparietal sulcus (IPS) bilaterally, but were not in schemas and words. Spatial directions were also decodable between all three formats. These results suggest that IPS plays a role in calculating spatial directions in visual scenes, but this neural circuitry may be bypassed when the spatial directions are presented as schemas or words.

**Significance Statement:** Human navigators encounter spatial directions in various formats: words (“turn left”), schematic signs (an arrow showing a left turn), and visual scenes (a road turning left). The brain must transform these spatial directions into a plan for action. Here, we investigate similarities and differences between neural representations of these formats. We found that bilateral intraparietal sulci represents spatial directions in visual scenes and across the three formats. We also found that participants respond quickest to schemas, then words, then images, suggesting that spatial directions in abstract formats are easier to interpret than concrete formats. These results support a model of spatial direction interpretation in which spatial directions are either computed for real world action, or computed for efficient visual comparison.

## Introduction

Humans fluently interpret spatial directions when navigating. But spatial directions are often presented in distinct representational formats – i.e., words, signs, and scenes – which have to be converted into a correct series of turns relative to one’s facing direction. While spatial directions conveyed through visual scenes are well-studied, little is known about how spatial directions are processed within and across these other frequently encountered formats. How does the human neural architecture for spatial cognition convert spatial maps and scenes into schematic maps and verbal directions?

Representational formats – visual scenes, schematic signs, and words – have distinct properties, which allow them to convey information differently. Visual scenes convey navigational information relatively directly – paths are visible – but contain irrelevant information (e.g., color, objects, context). Words, by contrast, categorize continuously varying turn angles and, by virtue of being symbolic, are related arbitrarily to the spatial directions conveyed. Schematic signs, or schemas, are exemplified by arrows in this investigation. Schemas are simplified visual representations of concepts (Talmy, 2000). Unlike visual scenes, schemas abstract over properties of spatial directions and omit those that are irrelevant. Unlike words, schemas maintain an iconic mapping between the spatial direction depicted and its representational format (i.e., a left-pointing arrow points to the left). Schemas may occupy a middle ground between images and words representing concepts (for actions, Watson, Cardillo, Bromberger, & Chatterjee, 2014; for prepositions, Amorapanth et al., 2012; c.f. Gilboa & Marlatte, 2017).

An intricate neural network interprets the spatial content of visual scenes. The occipital place area (OPA), parahippocampal place area (PPA), retrosplenial complex (RSC), and intraparietal sulcus (IPS) are implicated in different aspects of calculating spatial directions. The OPA codes egocentric spatial directions (anchored to one’s own body position) visible in visual scenes (Bonner and Epstein, 2017), whereas RSC codes allocentric spatial directions (anchored to properties of the environment) with respect to a known reference direction (i.e., a major axis of inside a building, (Marchette, Vass, Ryan, & Epstein, 2015; or north, Vass & Epstein, 2017). The PPA represents distinct spatial scenes and spatial directions relative to that scene (Epstein, 2008). Unlike OPA, PPA, and RSC, IPS codes egocentric spatial directions that can be either present in a scene or which were learned and then imagined. For example, Schindler & Bartels (2013) had participants memorize a circular array of objects and imagine movements with the same egocentric angle, but anchored to different objects (i.e., “face the lamp, point to the chair” and “face the chair, point to the vase” would both require a 60° clockwise rotation). Using multivoxel pattern analysis (MVPA), they showed that IPS exhibited similar patterns of activation for the same spatial direction. This work used visual scenes to encode spatial directions. Does IPS represent egocentric spatial directions from arrows (schematic depictions) and words similarly to visual scenes of egocentric spatial directions?

In the current work, we investigate how representational formats affect the behavioral and neural responses to spatial directions. Our broad hypothesis is that schemas and words elide the spatial processing required by visual scenes. If true, we predict evidence supporting two hypotheses: 1) schemas and words are processed more efficiently than scenes, and 2) visual scenes, but not schemas or words are processed in brain regions known to process spatial information. To test our first hypothesis, we predict that people will most quickly identify spatial directions depicted in words (“left” or “sharp right”) and schemas (arrows), compared to scenes (Google Maps images of roads). To test the second hypothesis, in an fMRI study using multivoxel pattern analysis (MVPA), we query the neural representations of spatial directions as a function of representational format. We expected visual scenes to be processed spatially and thus spatial directions would be decoded in IPS. We were agnostic if IPS would decode schemas and words, since these formats are not inherently spatial and need not be processed egocentrically. We also looked for cross-decoding between all three representational formats.

## Materials and Methods

### Participants

#### Norming Study

We recruited 42 participants (23 identifying as female) from Amazon Mechanical Turk. Two participants were removed for responding below chance. Of the remaining 40 participants (21 identifying as female, 1 did not report gender), 5 participants self-reported as Asian, 1 as African-American or Black, 2 as Hispanic, 1 as Other, and 29 as White. Two participants did not report ethnicity. Participants’ average age was 34.6 years (*SD* = 12.6). All but one participant reported speaking English as a first language.

#### Behavioral Study

We recruited 48 right-handed participants (27 identifying as female, 1 did not report gender) from a large urban university using an online recruitment tool specific to that university. 18 participants self-reported as Asian, 13 as African-American or Black, 1 as American Indian, 5 as Hispanic, 1 as Other, and 9 as Caucasian or White. One participant did not report ethnicity. Participants’ average age was 22.5 years (*SD* = 3.3). Participants reported speaking English as a first language.

#### fMRI Study

We recruited 22 right-handed participants from a large urban university using an online recruitment tool specific to that university. We excluded data from two participants because of motion. The resulting sample consisted of 20 participants (11 identifying as female). 4 participants self-reported as Asian, 4 as African-American or Black, and 12 as Caucasian or White. Participants’ average age was 21.4 years (*SD* = 2.7). Participants reported speaking English as a first language. Laterality quotient indicated participants were right-handed (*Min*. = 54.17, *M* = 80.63, *SD* = 16.41).

### Experimental materials

#### Stimuli

When given an open number of categories, people freely sort spatial directions into eight categories (Klippel and Montello, 2007). For the present study, we used seven of those eight categories: *ahead, left, right, sharp left, sharp right, slight left*, and *slight right*. We excluded *behind* because this direction would require the participant to imagine starting at the top of the image, rather than the bottom.

Spatial directions were depicted in three formats – words, schemas, and images. All stimuli were cropped to be 400×400 pixel squares. For each spatial direction, 24 words were created in Photoshop by modifying the size (small or large), font (Arial or Times New Roman), and color (blue, orange, pink, or purple). For each spatial direction, 24 schemas were created in Photoshop by modifying size (small or large), style (chevron or arrow), and color (blue, orange, pink, or purple). For the fMRI study, all 24 stimuli were used for each spatial direction. For the behavioral study, we psuedo-randomly chose three words and schemas to remove (retaining as close to the same number of colors, sizes, and fonts as possible across the directions), resulting in 21 exemplars per direction.

For each spatial direction, 28 images^1^ were created from Google Earth. Overhead satellite views were used to identify roads that turned in that direction, limiting the presence of cars, arrows on the road, and obscured view (like shadows or trees blocking the view). These 28 images were presented to 40 independent raters on Amazon Mechanical Turk, who answered a multiple choice question, selecting the spatial direction that best corresponded to the spatial direction depicted in the image. All raters rated all images. Across each image we compared the percentage of raters who selected the same direction we chose for each image as a measure of direction judgment reliability. We selected the top 30 raters on Mechanical Turk (eliminating the bottom nine participants who rated less than 75% of images correctly). For the fMRI study, we selected 24 images from each direction to match, as closely as possible, the agreement across spatial directions (overall agreement = 86.8%, *SD* = 11.6%). The images differed on overall agreement across spatial directions, *F*(6, 161) = 3.18, *p* = .006, ω^2^ = 0.07. Ahead images were the most agreed upon (*M* = 92.6%) and slight right images (*M* = 80.3%) were the least agreed upon. Across spatial directions, the difference in ratings was significant between the slight right images and the ahead images, *t*(46) = 2.62, *p* = .012, *d* = 0.77, and between the slight right images and the left images *t*(46) = 3.173, *p* = .003, *d* = 0.94. None of the other differences between pairs of spatial directions reached statistical significance. For the behavioral study, we used the most agreed upon 21 of these 24 images per spatial direction.

We created two versions of words and schemas. For the White background stimuli, words and schemas were displayed on a white square. For the Scrambled background stimuli, words and schemas were overlaid on phase-scrambled versions of the images. Two copies of each phase-scrambled image were used, to provide backgrounds for schemas and words. For Scrambled backgrounds, spatial directions and backgrounds were randomly paired for each participant. Sample stimuli (with phase-scrambled backgrounds) can be seen in Figure 1A.

**Figure 1.**
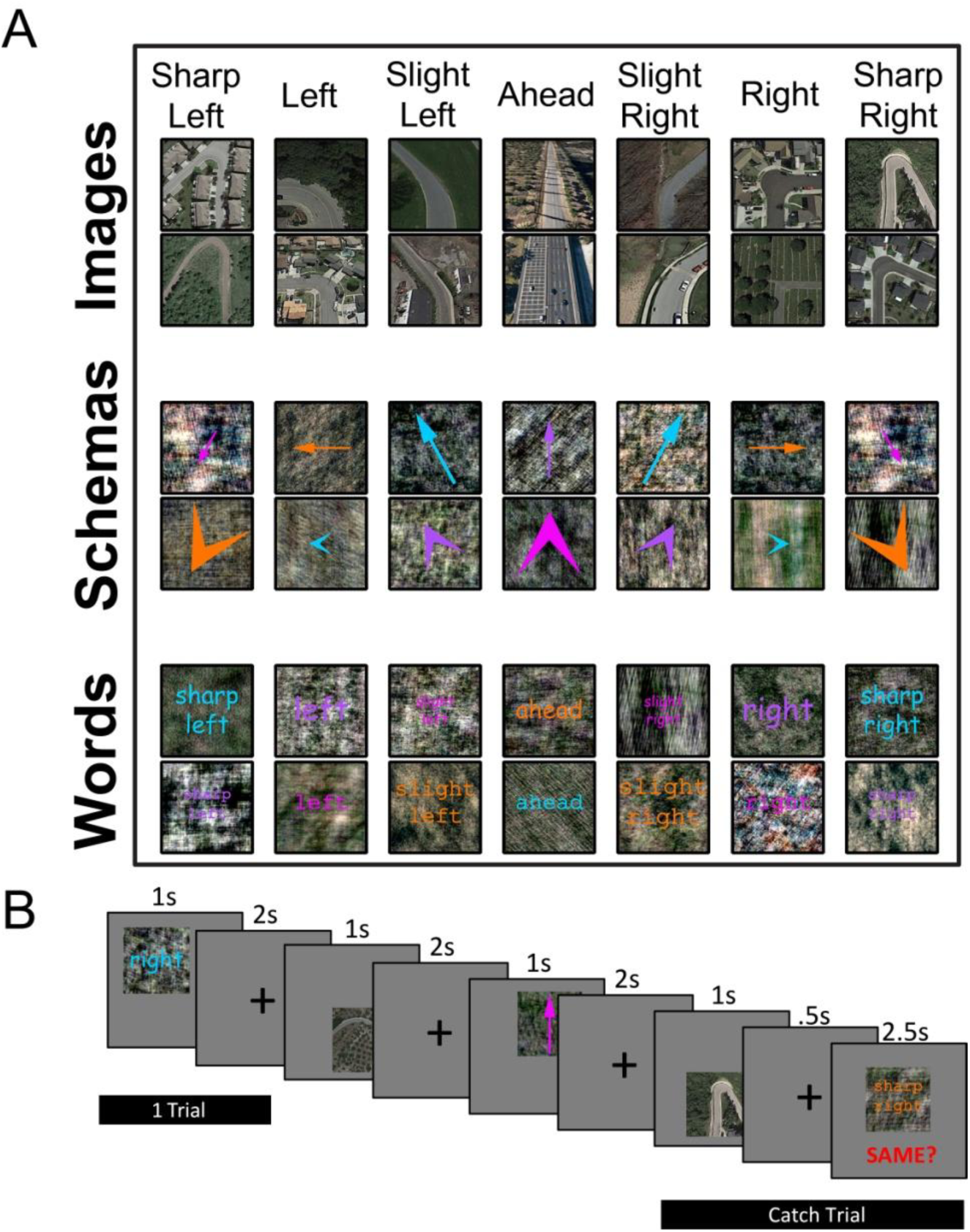
Stimuli and fMRI study paradigm. Sample stimuli from the behavioral and fMRI studies (A). Stimuli with phase shifted backgrounds are shown. A segment of the experimental paradigm shown to participants in the fMRI study (B). In the fMRI study, participants saw a spatial direction in one of the formats for 1 s, followed by a fixation cross for 2 s. During catch trials, the fixation cross instead disappeared after 500 ms, and a new spatial direction appeared, in any of the formats, with the word “Same” underneath in red letters. Participants pressed one button to indicate that the spatial direction on screen matched the one previously seen, and the other button to indicate that it did not match (buttons were counter-balanced across participants).

#### Self-report and debriefing

All participants in the fMRI and behavioral studies completed the Verbalizer-Visualizer Questionnaire (VVQ; Kirby, Moore, & Schofield, 1988) before the experimental task. Participants in the fMRI study also completed the Edinburgh handedness inventory (Oldfield, 1971) before the experimental task. After the experimental task, participants in the fMRI study completed a debriefing questionnaire, which asked how participants tried to remember the spatial directions and whether they felt it was difficult to switch between formats.

### Experimental Procedure (Behavioral Study)

To see how efficiently words, schemas, and images are encoded and translated across formats, participants performed a rolling one-back task. First, participants viewed instructions, which described the task and included samples of the seven spatial directions in all three formats. After a brief practice session, participants viewed spatial directions one at a time, responding by pressing one key to indicate that the current spatial direction was the same as the previously seen spatial direction, and another key to indicate that the current spatial direction was different from the previously seen direction. The spatial direction stayed on screen until the participant responded. Keys (‘F’ and ‘J’ on a standard keyboard) were counter-balanced across participants. Reaction time and accuracy were recorded. We generated a unique continuous carryover sequence for each participant (Aguirre, 2007) such that each spatial direction and format appeared before every other format and direction, including itself. This resulted in 441-trial sequences, of which approximately 1/7 were matches. Half of the participants performed the task with the White background spatial directions; the other half on the Scrambled background spatial directions. Because the first trial does not have a previous direction to compare, this trial was excluded from analysis.

### Experimental Procedure (fMRI Study)

To investigate the neural representations of spatial directions across and within formats, we presented spatial directions one at a time while the participant detected matches and non-matches in catch trials. Unlike the behavioral study, for the fMRI study we wanted to distinguish neural activation associated with individual spatial directions, rather than the comparison between one spatial direction and another. For that reason, participants only responded to occasional catch trials. The Scrambled spatial directions were used for the fMRI study because a computational model of early visual cortex processing (the Gist model: Oliva & Torralba, 2006) could not cross-decode spatial direction across formats when scrambled backgrounds were used, but could when white backgrounds were used. Using the scrambled backgrounds in the fMRI study reduced the likelihood of decoding spatial directions across formats because early visual cortex might be sensitive to low level visual properties that could, for example, distinguish schemas and images.

Continuous carryover sequences (Aguirre, 2007) were generated with 24 conditions - seven spatial directions in each of the three formats made up 21 conditions; two catch trials (one match and one non-match); and one null condition. The resulting sequence consisted of 601 trials (504 spatial directions trials, 48 catch trials, 48 null trials, and the first trial repeated at the end). Except for the catch trials, which could consist of any stimulus, exemplars were only presented once per participant. A schematic of the trial structure can be seen in Figure 1B.

For spatial direction trials, participants viewed spatial directions one at a time, presented on screen for 1000 ms with a 2000 ms inter-stimulus interval consisting of a fixation cross. Participants were instructed to attend to the spatial direction for each trial. On catch trials, a subsequent spatial direction appeared with the word “Same?” underneath in large red letters. For these trials, participants pressed one key to indicate that the spatial direction was the same as the one they just saw, and another key to indicate that the spatial direction was different. Keys (the leftmost and rightmost buttons of a four-button MRI-compatible button box) were counter-balanced across participants.

Catch trials consisted of 1000 ms stimulus presentation, followed by 500 ms fixation, then 4500 ms of the catch stimulus. The catch stimulus was randomly chosen each time, with the constraint that it could not be the exact same stimulus. If the catch trial was a match trial, the spatial direction had to be the same. If the catch trial was a non-match trial, the spatial direction was randomly chosen from all the other spatial directions. Catch stimuli could be any format. Null trials consisted of a fixation cross, presented for double the normal trial length, 6000 ms (Aguirre, 2007).

The experimental session was divided into 6 runs. The runs were 100 trials each, except for the last, which was 101 trials. The second through sixth runs began with the last five trials of the previous run to re-instate the continuous carryover sequence. These overlap trials, as well as the catch trials, and null trials, were not analyzed. Because runs contained between 6-9 catch trials, which were 6000 ms, the runs varied slightly in length, but were approximately 5 min, 50 s. Because of this variation, the scanner collected functional data for 6 min, 12 s. Additional volumes collected after the stimuli for those trials were finished were discarded. Reaction time and accuracy were recorded. After each run, the participant received feedback on his performance (e.g., “You got 6 out of 8 correct.”).

### MRI Acquisition

Scanning was performed at the Hospital of the University of Pennsylvania using a 3T Siemens Trio scanner equipped with a 64-channel head coil. High-resolution T1-weighted images for anatomical localization were acquired using a three-dimensional magnetization-prepared rapid acquisition gradient echo pulse sequence [repetition time (TR), 1850 ms; echo time (TE), 3.91 ms; inversion time, 1100 ms; voxel size, 0.9 × 0.9 × 1 mm; matrix size, 240 × 256 × 160]. T2*-weighted images sensitive to blood oxygenation level-dependent (BOLD) contrasts were acquired using a multiband gradient echo echoplanar pulse sequence (TR, 3000 ms; TE, 30 ms; flip angle, 90°; voxel size, 2 × 2 × 2 mm; field of view, 192; matrix size, 96 × 96 × 80; acceleration factor, 2.). Visual stimuli were displayed by rear-projecting them onto a Mylar screen at 1024 × 768 pixel resolution with an Epson 8100 3-LCD projector equipped with a Buhl long-throw lens. Participants viewed the stimuli through a mirror attached to the head coil.

Functional images were corrected for differences in slice timing using FSL slice-time correction and providing the interleaved slice time order. Images were then realigned to the first volume of the scan run, and subsequent analyses were performed within the participants’ own space. Motion correction was performed using MCFLIRT (Jenkinson et al., 2002), but motion outliers were also removed using the Artifact Detection Toolbox (http://www.nitrc.org/projects/artifact_detect).

For two participants, data from two runs were discarded. For one participant, data were excluded because the scanning computer crashed during the final run. For a second participant, data were excluded because performance on the behavioral task was below 50% (chance) for the final run. All other runs for all other participants exceeded 63% correct.

### Multivoxel pattern analysis

To test whether regions of the brain encoded information about spatial direction, we calculated, within each participant, the similarities across scan runs between the multivoxel activity patterns elicited by each spatial direction in each format. If a region contains information about spatial direction, then patterns corresponding to the same direction in different scan runs should be more similar than patterns corresponding to different directions (Haxby et al., 2001). Moreover, if this effect is observed for patterns elicited by stimuli of different formats (i.e., word-schema), then the spatial direction code generalizes across formats.

To define activity patterns, we used general linear models (GLMs), implemented in FSL (Jenkinson et al., 2012), to estimate the response of each voxel to each stimulus condition (three formats for each of seven spatial directions) in each scan run. Each runwise GLM included one regressor for each spatial direction in each format (21 total), regressors for motion parameters, and nuisance regressors to exclude outlier volumes discovered using the Artifact Detection Toolbox (http://www.nitrc.org/projects/artifact_detect/). Additional nuisance regressors removed catch trials and the reinstatement trials which began runs 2-5. High-pass filters were used to remove low temporal frequencies before fitting the GLM, and the first three volumes of each run were discarded to ensure data quality. Multivoxel patterns were created by concatenating the estimated responses across all voxels within either the region of interest or the searchlight sphere. These patterns were then averaged across the first three runs, and then across the second three runs. For the two participants for whom the final run was discarded, the last two runs were averaged together.

To determine similarities between activity patterns, we calculated Kendall’s τ_A_ correlations (Nili et al., 2014) between patterns in the first half and second half scan runs. Before this computation, we removed the cocktail mean (the average neural activity pattern across all conditions; Vass and Epstein, 2013) within each format and within each run separately. This approach normalizes activity patterns across conditions. The pattern of results was unchanged when the cocktail mean was not removed.

We then performed representational similarity analyses by comparing the correlations between the neural signal across conditions to a theoretical representational dissimilarity matrix (RDM), which specified how the data would look if a hypothesis were true. This comparison occurred in one of two ways. 1) If the theoretical RDM was continuous, we correlated the neural RDM with the theoretical RDM. 2) If the theoretical RDM was binary, we obtained a discrimination index by averaging a subset of correlations from the neural RDM (e.g., different direction correlations) and subtracting that from the average of another subset (e.g., same direction correlations).

### Searchlight analysis

To test for format decoding across the brain, we implemented a whole-brain searchlight analysis (Kriegeskorte et al., 2006) in which we centered a spherical ROI (radius, 5 mm) around every voxel of the brain, calculated the spatial direction correlation within this spherical neighborhood using the method described above, and assigned the resulting value to the central voxel. Searchlight maps from individual participants were then aligned to the Montreal Neurological Institute (MNI) template with a linear transformation and submitted to a second-level random-effects analysis to test the reliability of discrimination across participants. To find the true type I error rate, we performed Monte Carlo simulations that permuted the sign of the whole-brain maps from individual participants (Nichols and Holmes, 2002; Winkler et al., 2014). We performed this procedure 1,000 times across the whole brain. Voxels were considered significant if they exceeded the *t*-statistic of the top 5% of permutations. The mean chance correlation was 0.

### Regions of interest

#### Scene-selective regions

We identified scene-selective regions of interest (ROIs), Figure 2A-B. These ROIs were defined for each participant individually using a univariate contrast of images>words+schemas, and a group-based anatomical constraint of scene-selective activation derived from a large number (42) of participants from a previous study (Julian et al., 2012). Specifically, each ROI was defined as the top 100 voxels in each hemisphere that responded more to images than to words+schemas and fell within the group-parcel mask for the ROI. To avoid double-dipping, we defined the ROI using the image>word+schema contrast for one run, then performed the MVPA analysis as described above on the remaining runs. This method ensures that all scene-selective ROIs could be defined in both hemispheres in every participant and that all ROIs contain the same number of voxels, thus facilitating comparisons between regions.

**Figure 2.**
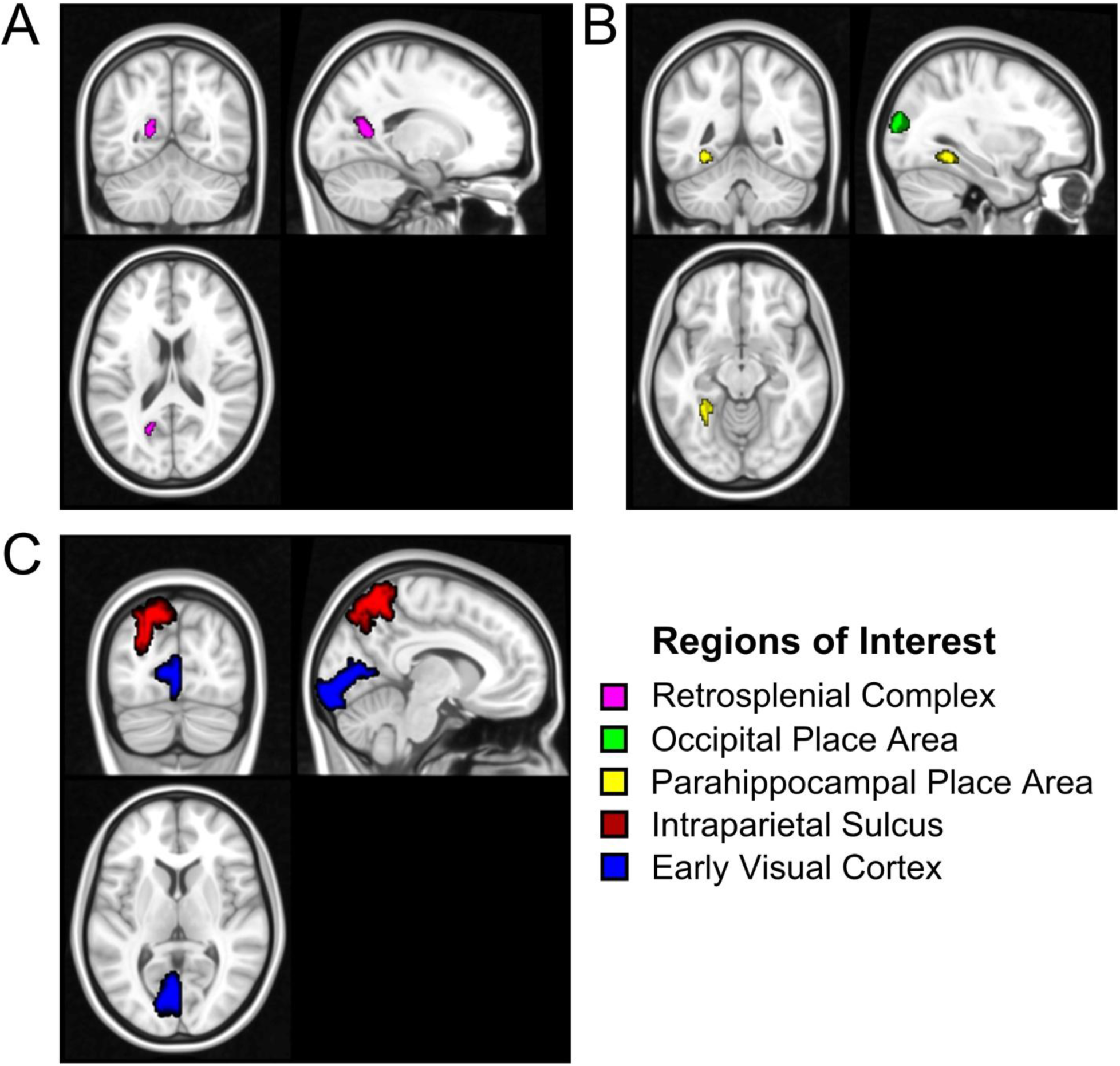
Regions of Interest. The five regions of interest (ROIs) used in the fMRI analyses, shown for illustrative purposes (data analysis proceeded as described in the Methods). ROIs are also displayed in the left hemisphere only, for ease of viewing, but were analyzed bilaterally. The scene regions (Parahippocampal Place Area, PPA, yellow; Occipital Place Area, OPA, green; Retrosplenial Cortex, magenta) are displayed as the top 100 voxels of the group averaged t-statistic for the contrast of images - mean(words + schemas), and constrained by the anatomical parcels from Julian et al. (2012). Intraparietal sulcus (IPS, red) and early visual cortex (EVC, blue) were defined anatomically by the parcels from Wang et al. (2014). All ROIs are displayed on the standard MNI map.

#### Visual and Parietal regions

We defined early visual cortex (EVC) and intraparietal sulcus (IPS) using the probabilistic atlas from Wang and colleagues (Wang et al., 2015) Figure 2C. These parcels were registered to participants’-own-space and voxels were extracted. The MVPA analysis was then performed as described above using all data. We analyzed all IPS regions in one combined ROI (the union of all voxels from the Wang et al IPS parcels) without further selection of voxels from functional comparisons.

### Experimental Design and Statistical Analysis

We conducted 2 experimental studies. The sample size for the behavioral study was selected based on a power analysis from a smaller pilot study with 15 participants. The behavioral study and reported analyses were preregistered on Open Science Framework (https://osf.io/5dk37/), but the code used to analyze the data was altered, because of software bugs, which were unknown at the time of the original registration. Within- and between-subject factors and materials for that study can be found on Open Science Framework, and in the method section above.

The sample size for the imaging study was based on previous similar studies (Schindler and Bartels, 2013; Marchette et al., 2015) that examined within-subject differences in MVPA of the BOLD fMRI signal. Although we looked at individual differences in an exploratory fashion, we interpret these results with caution. Details and important parameters for the imaging study can be found in the Method section.

Across both studies, where appropriate, we corrected for multiple comparisons and report in the text how these determinations were made.

## Results

### Behavioral Study

Accuracy on the rolling one-back task was high (*M* = 93.7%, *SD* = 21.57, Range = [75.0% - 99.8%]). Including incorrect trials, reaction time overall was *M* = 1.31s, *SD* = 0.34s, Range = [0.70s – 2.14s]. We excluded data from one participant because of low accuracy (53.9%) and fast reaction time (0.24s) compared to data from the rest of the sample.

Participants did not differ in accuracy when responding to schemas (*M* = 94.7%, *SD* = 4.2%), and words (*M* = 94.4%, *SD* = 4.2%), *t*(46) = 1.21, *p* = .23, d = 0.14, but were more accurate responding to schemas compared to images (*M* = 91.3%, *SD* = 7.2%), *t*(46) = 5.26, *p* = .0000004, *d* =0.97. Participants were also more accurate for words compared to images, *t*(46) = 4.68, *p* = .000003, *d* = 0.88. This result shows that, compared to schemas and words, the spatial directions in the images were more difficult to identify (i.e., it was possible to interpret a slight right turn as a right or an ahead). To avoid this confound and a speed-accuracy tradeoff, we excluded incorrect trials and only analyzed reaction times for correct trials across formats.

#### Schemas are processed more quickly than images or words

In addition to excluding correct trials, we excluded trials for which the participant responded especially slowly – greater than two standard deviations above his/her mean reaction time. We also excluded trials for which the answer was “same,” because these trials occurred relatively infrequently and could be considered oddball trials. They also required a different response than the other trials. All further analyses exclude trials as described above.

Figure 3 displays the main reaction time results for the behavioral study. Reaction time for schemas was quicker (*M* = 1.13s, *SD* = 0.28s) than for images (*M* = 1.27s, *SD* = 0.35s), *t*(46) = 7.40, *p* = .000000002, *d* = 1.19, and words (*M* = 1.18s, *SD* = 0.26s), *t*(46) = 3.09, *p* = .003, *d* = 0.49. Reaction time for words was also quicker than for images, *t*(46) = 4.38, *p* = .00007, *d* = 0.97.

**Figure 3.**
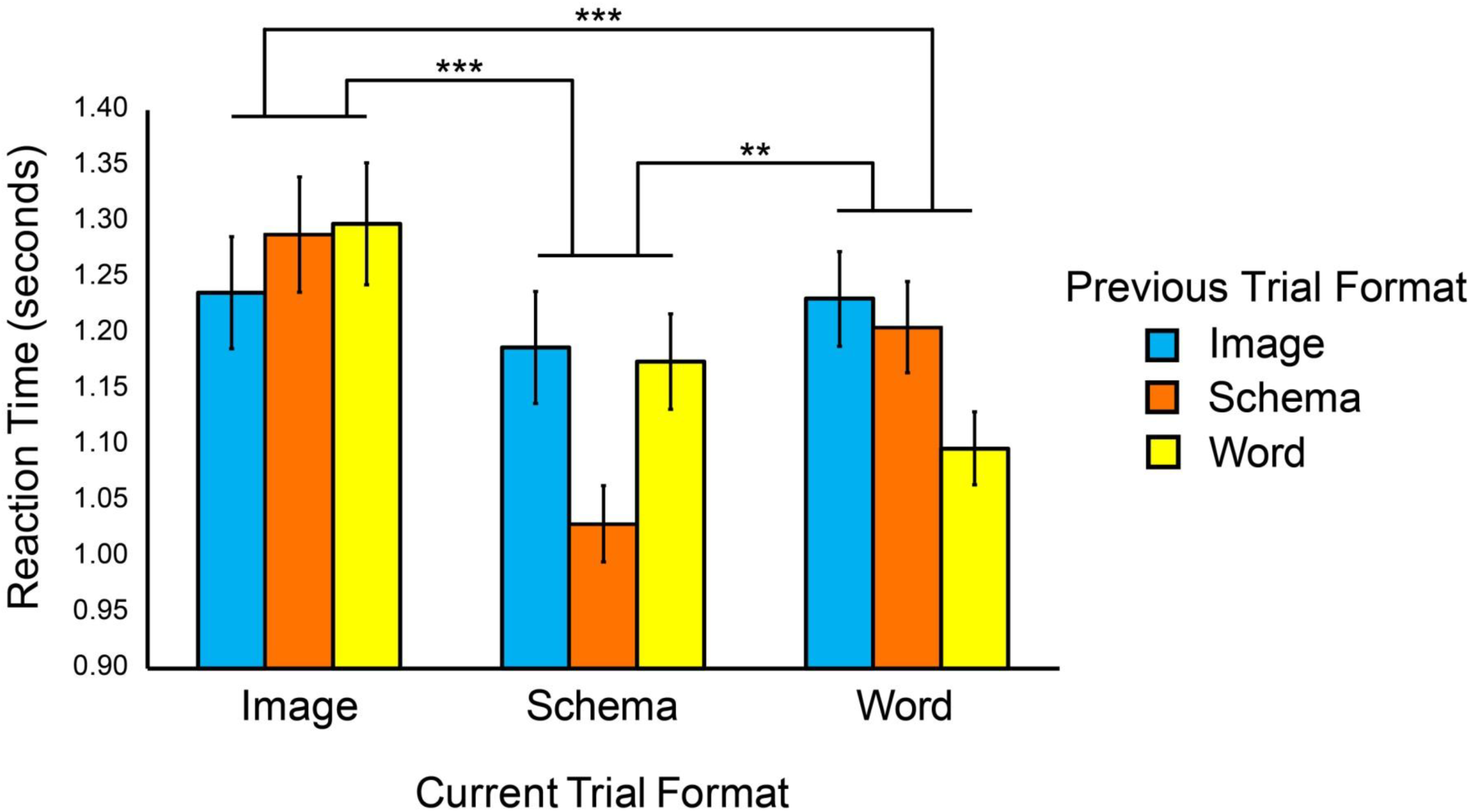
Results from the behavioral study. Response times were fastest overall for schemas and words. Schemas also showed the largest within-format effect. That is, participants were faster to respond when a schema came after a schema compared to word-word or image-image.

#### Same-format advantage

Comparing spatial directions was faster when the preceding stimulus was in the same format. We calculated the average reaction time for each current format (the trial for which a response is generated) separately based on whether the previous trial was the same or a different format. The same-format advantage is operationalized as the difference in reaction time between same-format-preceding trials and different-format-preceding trials. Higher numbers indicate faster responses for same-format comparisons than different-format comparisons. Images (*M* = 0.06s, *SD* = 0.16s), one-sample *t*(46) = 2.47, *p* = .017, *d* = 0.38, schemas (*M* = 0.15s, *SD* = 0.15s), one-sample *t*(46) = 6.94, *p* = .00000001, *d* = 1.00, and words (*M* = 0.12s, *SD* = 0.13s), one-sample *t*(46) = 6.38, *p* = .00000008, *d* = 0.92, all showed significant same-format advantages. Comparing the same-format advantage between images, schemas, and words revealed that schemas showed a larger same-format advantage than images, *t*(46) = 3.43, *p* = .001, *d* = 0.50, and a marginally larger advantage than words, *t*(46) = 1.86, *p* = .07, *d* = 0.34. Words showed a significantly larger same-format advantage than images, *t*(46) = 2.35, *p* = .02, *d* = 0.28. Comparing spatial directions was always faster when these comparisons were within format, but schemas showed this effect most strongly.

#### Phase-scrambled backgrounds did not show behavioral effects

As reported in the Method section, the Gist model could not decode spatial directions across formats when scrambled backgrounds were used for schemas and words, but could do so with white backgrounds. Despite this finding, we did not find behavioral effects based on background. Reaction time (on all trials) was similar for White (*M* = 1.28s, *SD* = 0.32s) and Scrambled backgrounds (*M* = 1.33s, *SD* = 0.36s), *t*(45) = 0.43, *p* = .67, *d* = 0.15. Accuracy was similar for White (*M* = 93.1%, *SD* = 5.50%) and Scrambled backgrounds (*M* = 94.2%, *SD* = 4.30%), *t*(45) = 0.76, *p* = .45, *d* = 0.22. In addition, none of the above analyses interacted with the background condition.

#### Reaction time correlates with egocentric not visual angular distance between trials

We instructed participants to imagine the directions as egocentric, with respect to their own body position, but wondered whether reaction time data were consistent with participants following this instruction. Thus, we calculated the angular distance between each pair of trials in two ways. The visual angle was calculated as the absolute value of the angular distance between the current and previous trials. The egocentric angle was calculated similarly, except that all angular distances were calculated as if sharp right and sharp left were maximally far apart. We called this the egocentric angle because relative to one’s facing direction, the head cannot rotate behind the body, thus this angle calculation preserves egocentric validity. For example, the angular distance between sharp right and sharp left was 90° for visual angle, but 270° for egocentric angle. To determine whether there was a significant correlation within participants, we calculated the Pearson’s correlation between each participant’s reaction time on that trial with the visual and egocentric angular distance between that trial and the previous trial. We then conducted one-sample *t*-tests to determine if there was a significant correlation in our sample, and within-subject *t*-tests to compare correlations. We found that egocentric angles correlated with reaction time positively (*M* = .040, *SD* = .065), *t*(46) = 4.21, *p* = .0001, *d* = 0.61, but visual angles did not (*M* = .0074, *SD* = .064), *t*(46) = 0.79, *p* = .43, *d* = 0.12. These patterns significantly differed from each other, *t*(46) = 2.75, *p* = .009, *d* = 0.82. This pattern of results was obtained within each format separately, and angular distance correlation did not interact with format. This pattern of results reveals that participants interpreted spatial directions egocentrically because longer reaction times were associated with larger egocentric but not visual angle distances.

#### Individual differences in cognitive style did not correlate with reaction time

No measures of reaction time correlated with either measure of the Verbalizer-Visualizer Questionnaire, all *p*’s > .08.

### fMRI Study

#### Behavioral performance during the fMRI task

Responses to the catch trials during the fMRI task were accurate (*M* = 89.9%, *SD* = 6.74%). Behavioral responses during one run for one participant fell below chance (43%, 3/7 correct). fMRI data for that run for that participant were excluded.

#### Spatial direction decoding

##### Within-format decoding of spatial direction in ROIs

Figure 4A displays the within-format contrasts used to calculate whether spatial directions were decoded in each ROI. The whole grid represents a theoretical RDM of three separate contrasts: Same minus different spatial direction within each format. These contrasts were performed on each participant’s neural RDM, calculated as described in the Method section by correlating the averaged parameter estimates for each trial type (e.g., a slight right word, or a sharp left schema) separately for the first and second half of each participant’s runs. Separately for each format, grey squares were subtracted from colored squares. White squares were omitted.

**Figure 4.**
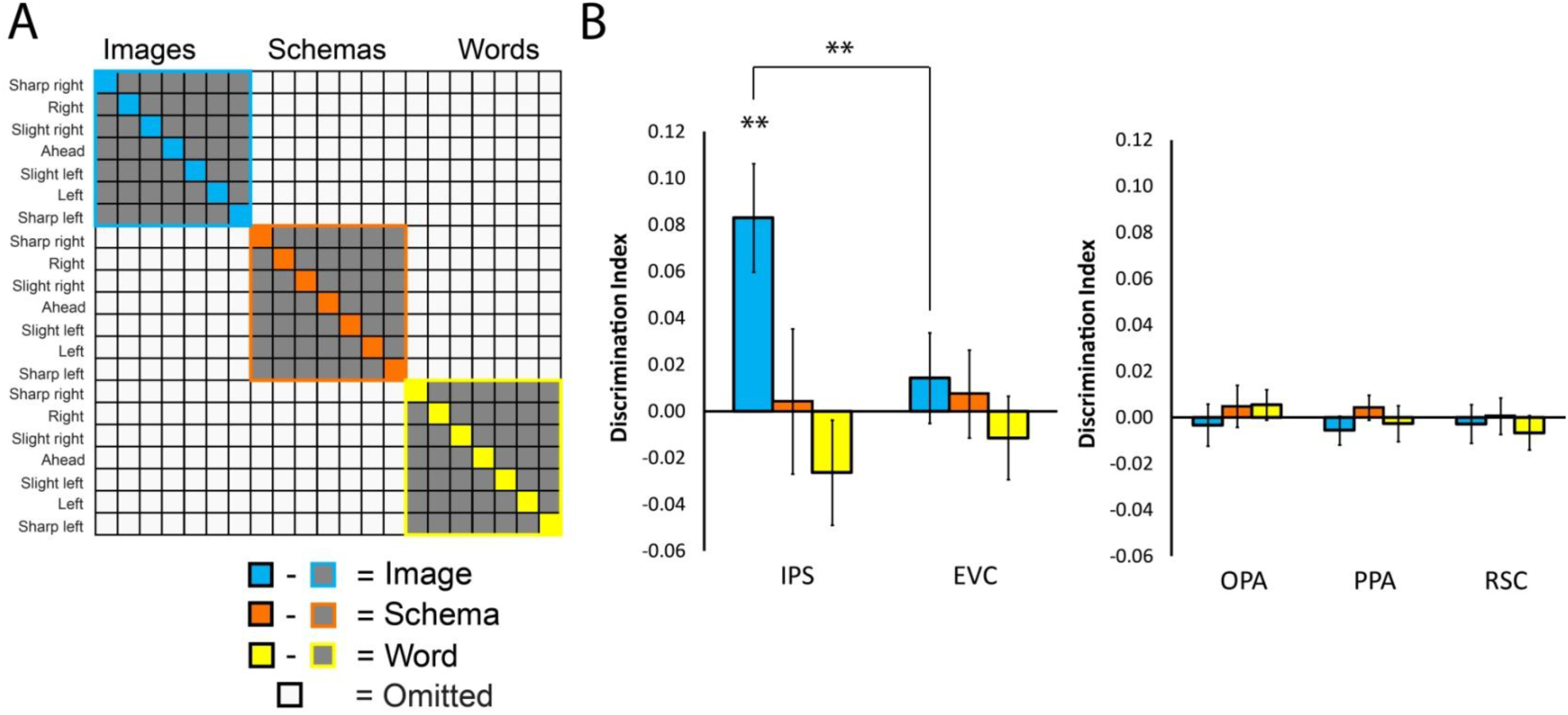
Within-format decoding of spatial direction. The theoretical RDM (A) was compared to the neural RDM from five ROIs: the intraparietal sulcus (IPS), early visual cortex (EVC), and visual scene regions: occipital place area (OPA), parahippocampal place area (PPA), and retrosplenial complex (RSC). The IPS decoded spatial directions within images significantly greater than EVC. Within-format decoding was not significant in IPS or EVC for either schemas or words. Visual scene regions did not decode spatial direction within any of the three formats.

Within-format results for each of the ROIs are displayed in Figure 4B. Spatial direction was decoded within images in IPS (*M* = 0.08, *SD* = 0.10), one-sample *t*(19) = 3.56, *p* = .002, *d* = 0.80. EVC did not decode spatial direction within images, (*M* = 0.01, *SD* = 0.09), one-sample *t*(19) = 0.73, *p* = .47, *d* = 0.10. IPS decoded spatial direction within images significantly more than EVC, *t*(19) = 2.97, *p* = .0078, *d* =0.69. Scene regions did not decode spatial directions within images (*M*_OPA_ = -0.003, *SD*_OPA_ = 0.04, *t*_OPA_(19) = 0.36, *p* = .72, *d* = -0.08; *M*_PPA_ = -0.006, *SD*_PPA_ = 0.03, *t*_PPA_(19) = 0.90, *p* = .38, *d* = -0.20; *M*_RSC_ = -0.003, *SD*_RSC_ = 0.04, *t*_RSC_(19) = 0.34, *p* = .74, *d* = -0.08).

Spatial directions were not decoded for schemas or for words in any of the ROIs (all *p*’s > .26).

##### Cross-format decoding of spatial direction in ROIs

We also wished to learn if spatial directions could be decoded independently of the visual properties of individual formats. A brain region would show evidence of cross-format decoding of spatial direction if the correlation between the same spatial direction, presented in different formats, exceeded the correlation between different spatial directions, presented in different formats. Figure 5A displays the cross-format decoding theoretical RDM. Grey squares (different direction, different format) are subtracted from black squares (same direction, different format) to yield the degree of generalization. White squares are omitted.

**Figure 5.**
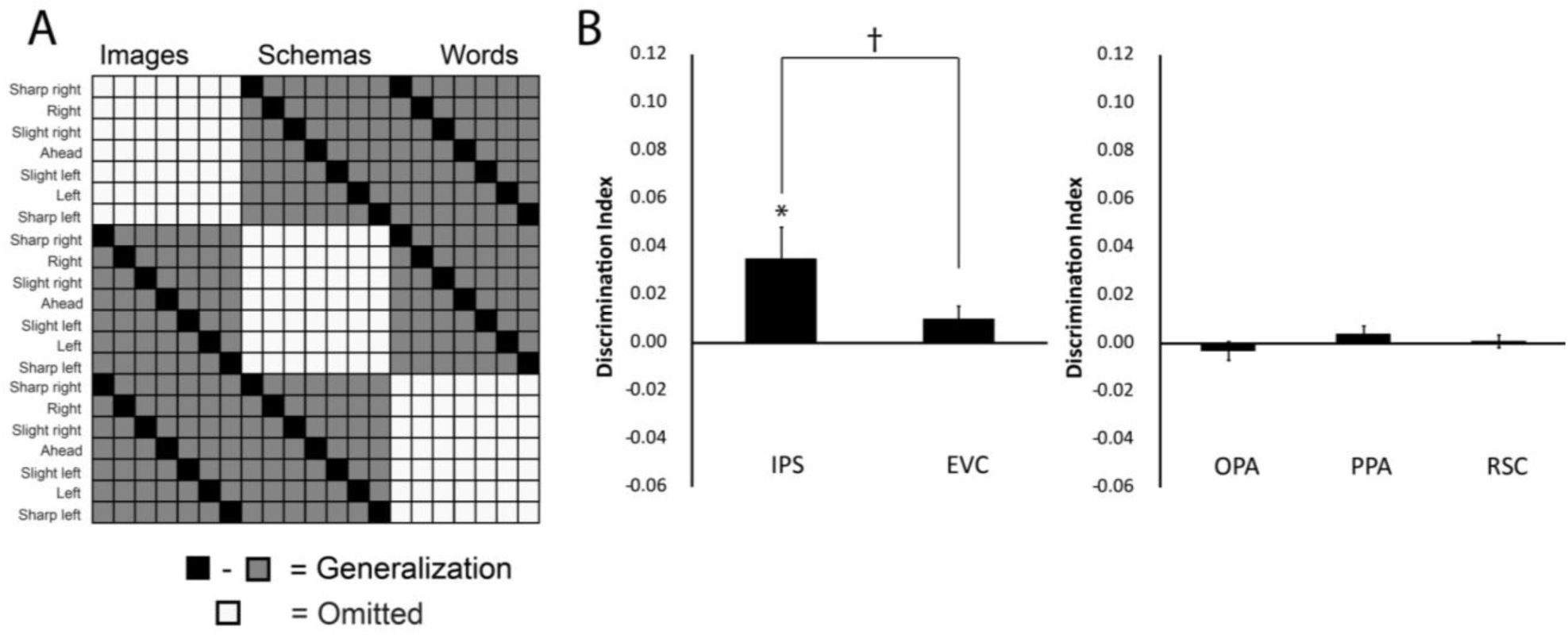
Cross-format decoding of spatial direction. The theoretical RDM (A) was compared to the neural RDM from five ROIs: the intraparietal sulcus (IPS), early visual cortex (EVC), and visual scene regions: occipital place area (OPA), parahippocampal place area (PPA), and retrosplenial complex (RSC). The IPS decoded spatial directions across all three formats, but only marginally greater than EVC. Cross-format decoding was not significant in IPS or EVC for either schemas or words. Visual scene regions did not decode spatial direction across formats.

Results for each of the ROIs are displayed in Figure 5B. Cross-format spatial directions were decoded in IPS (*M* = 0.04, *SD* = 0.06), one-sample *t*(19) = 2.64, *p* = .0128, *d* = 0.67. There was marginally significant cross-format decoding in EVC (*M* = 0.01, *SD* = 0.02), one-sample *t*(19) = 1.80, *p* = .087, *d* = 0.50. IPS decoded spatial directions across formats marginally more than EVC, *t*(19) = 2.09, *p* = .051, *d* = 0.66. The scene regions did not decode spatial directions across formats (*M*_OPA_ = -0.003, *SD*_OPA_ = 0.02, *t*_OPA_(19) = 0.79, *p* = .44, *d* = -0.15; *M*_PPA_ = 0.004, *SD*_PPA_ = 0.015, *t*_PPA_(19) = 1.21, *p* = .24, *d* = 0.27; *M*_RSC_ = 0.0009, *SD*_RSC_ = 0.01, *t*_RSC_(19) = 0.34, *p* = .74, *d* = 0.09). These data reveal that IPS contain cross-format representations of spatial direction, but EVC and scene regions do not.

Within IPS we wanted to know whether cross-format decoding of spatial direction was driven by particular pairs of formats. For example, it is possible that spatial direction decoding was high between images and schemas, but comparatively lower between images and words. To investigate this, we conducted follow-up contrasts between each pair of formats similar to the omnibus test above (e.g., same direction, different format minus different direction, different format for images versus schemas). These follow-up contrasts revealed significant schema-word decoding (*M* = 0.036, *SD* = 0.07), one-sample *t*(19) = 2.23, *p* = .04, *d* = 0.51 and marginally significant image-schema decoding (*M* = 0.024, *SD* = 0.05), one-sample *t*(19) = 1.99, *p* = .06, *d* = 0.48. Image-word decoding was not significant (*M* = 0.008, *SD* = 0.07), one-sample *t*(19) = 0.52, *p* = .61, *d* = 0.11. These follow-up contrasts were not significantly different from each other (all pairwise *p*’s > .25). This pattern of results suggest that schemas may occupy an intermediary role, sharing neural responses in IPS with images and words respectively in a way not seen with images and words.

##### Spatial direction decoding in searchlights

Although we had specific predictions about regions of the brain that might decode spatial directions, we also conducted exploratory analyses to assess within- and cross-format spatial direction decoding at the whole-brain level. We did so to see whether any regions of the brain outside IPS decoded spatial directions in words or schemas. None of these analyses survived correction for multiple comparisons. So we report our observations at a lower threshold and advise caution in interpretation. We report results that exceed a lower threshold (p < .0005, uncorrected). Within-format decoding of images occurred in left posterior parietal cortex, extending into left medial parietal cortex – a region consistent with our IPS ROI, as well as a left lateral frontal region. Within-format decoding of schemas occurred in left premotor cortex, and within-format decoding of words occurred in a small region in the brain stem. Cross-format decoding revealed a small region near left visual area MT.

#### Spatial direction similarity analysis

The preceding analyses reveal that IPS can distinguish between the seven spatial directions within images, and across formats. There are two possible ways IPS could do this. IPS could be creating seven arbitrary and ad hoc categories for each spatial direction, which could allow any type of information to be decoded. If this interpretation is correct, the IPS’ role in spatial direction coding would be that it is creating a problem space onto which any possible stimulus categories could be mapped. For example, if the task were to sort stimuli based on seven colors, IPS would create seven color categories, which would be most similar to themselves (e.g., red is most similar to red), and different from all others. On the other hand, IPS could be involved because it helps distinguish spatial directions, specifically. If this interpretation is correct, the IPS’ role in spatial direction coding would be that it constructs a spatial representation of the possible directions. A counter-example for color would be that IPS contains a color-wheel representation. To distinguish which of these possibilities is correct, we can analyze off-diagonal spatial direction similarity. We would expect categories of turns (e.g., left to slight left) to be more similar to each other than to more distant turns (e.g., left to sharp right). We created a new theoretical RDM in which all left turns (sharp left, left, and slight left) were similar to each other, and dissimilar to all right turns (and vice versa for right to left turns). Ahead directions were coded as dissimilar from everything else. We excluded the diagonal to ensure that these results are not recapitulations of the spatial direction decoding analyses above. That is, this analysis captures similarity among non-identical spatial directions to show that IPS neural patterns contain spatial information (not arbitrary category information).

We found that the neural pattern of activity in IPS in response to images correlated more strongly between left turns than across left and right turns (*M* = 0.036, *SD* = 0.063), *t*(19) = 2.59, *p* = .018, *d* = 0.57. This pattern was not the case for schemas, (*M* = -0.018, *SD* = 0.086), *t*(19) = 0.09, *p* = .93, *d* = -0.21, nor for words, (*M* = -0.017, *SD* = 0.075), *t*(19) = 1.02, *p* = .32, *d* = -0.23, nor across formats, (*M* = 0.009, *SD* = 0.037), *t*(19) = 1.10, *p* = .29, *d* = 0.24. This result provides evidence that images were represented spatially, by distinguishing left from right turns, in IPS, and not as seven arbitrary and ad hoc categories. Although this analysis shows that IPS codes spatial content, the theoretical RDM we chose was not the only possible one. We also conducted a representational similarity analysis wherein we correlated the neural RDM with a spatial direction model where similarity linearly decreased as a function of spatial angle, but this analysis did not achieve statistical significance. We thus interpret this result as evidence of spatial content in IPS, but do not feel strongly that the representation is categorical (i.e., all lefts are more similar to each other than to rights).

#### Format decoding

In the following analyses, we removed the cocktail mean within run, across all formats.

##### Format decoding in ROIs

In addition to direction coding, we wanted to determine whether the format of stimuli was represented in these ROIs. The theoretical RDM for this contrast is presented in Figure 6A. For this analysis, we excluded correlations between stimuli that were the same direction and the same format (white squares in Figure 6A). To decode format, a region would show higher correlations between stimuli that were the same format compared to stimuli that were different formats (black squares minus grey squares in Figure 6A).

**Figure 6.**
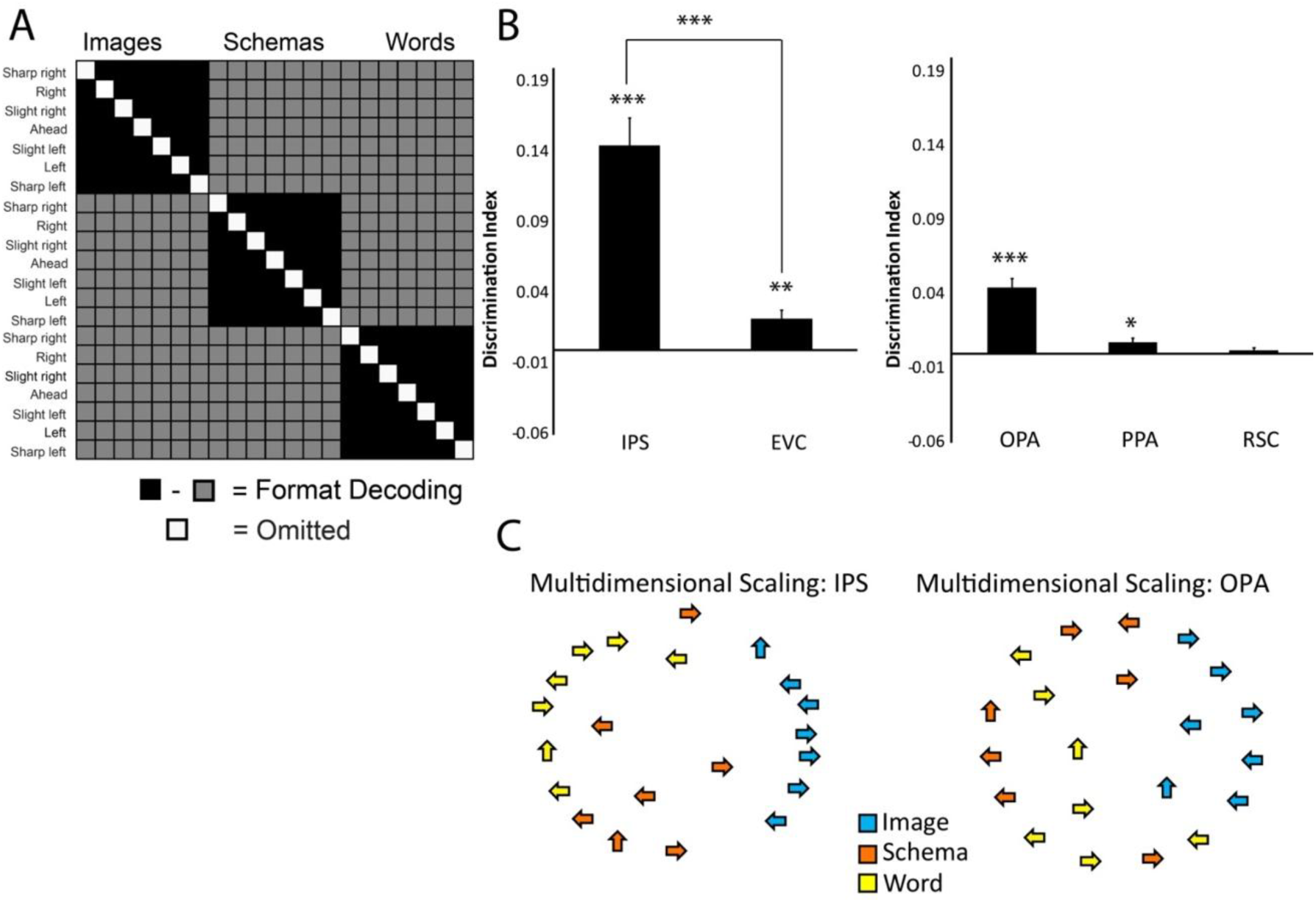
Decoding of format in ROIs. The theoretical RDM (A) was compared to the neural RDM from five ROIs: the intraparietal sulcus (IPS), early visual cortex (EVC), and visual scene regions - occipital place area (OPA), parahippocampal place area (PPA), and retrosplenial complex (RSC). All regions significantly decoded the format of the representation, except for RSC. Multidimensional scaling plots (C) reveal that IPS separates all three formats whereas OPA distinguishes images from the other two. Arrows depict that categorical spatial directions (right, slight right, and sharp right collapsed as right arrows; left, slight left, and sharp left collapsed as left arrows; ahead as an up arrow).

Results from the omnibus format decoding contrast can be seen in Figure 6B. Format could be decoded in IPS (*M* = 0.14, *SD* = 0.09), one-sample *t*(19) = 7.25, *p* = .0000007, *d* = 1.56, and EVC (*M* = 0.02, *SD* = 0.03), one-sample *t*(19) = 3.58, *p* = .002, *d* = 0.67, although format decoding was significantly higher in IPS than EVC t(19) = 6.39, *p* = .000004, *d* = 1.63. OPA (*M* = 0.04, *SD* = 0.03), one-sample *t*(19) = 7.07, *p* = .000001, *d* = 1.33, and PPA (*M* = 0.008, *SD* = 0.01), one-sample *t*(19) = 2.54, *p* = .02, *d* = 0.80, also decoded format, although RSC did not (*M* = 0.003, *SD* = 0.008), one-sample *t*(19) = 1.39, *p* = .18, *d* = 0.38.

We wanted to know whether the regions that significantly decoded format generally (IPS, EVC, OPA, and PPA) could decode pairwise formats. We thus looked at schema-word, schema-image, and image-word decoding separately for each ROI. See Table 1 for the complete results. In sum, pairwise formats could be decoded to some extent in each ROI except RSC. In IPS and OPA, all three pairs of formats could be distinguished, whereas PPA predominantly dissociated images from the schemas and words.

**Table 1.** IPS (Intraparietal Sulcus). EVC (Early Visual Cortex). OPA (Occipital Place Area). PPA (Parahippocampal Place Area).

| Brain region | Schema-Word |  |  |
| --- | --- | --- | --- |
|  | Mean (Standard deviation) | *p*-value | Effect Size (*d*) |
| IPS | .034(.037) | 0.0007 | 0.92 |
| EVC | .025(.035) | 0.0057 | 0.71 |
| OPA | .006(.035) | 0.017 | 0.17 |
| PPA | -.0006(.009) | 0.77 | 0.07 |
|  | Schema-Image |  |  |
|  | Mean (Standard deviation) | *p*-value | Effect Size (*d*) |
| IPS | .19(.127) | 0.0000008 | 1.5 |
| EVC | .027(.039) | 0.0057 | 0.69 |
| OPA | .06(.04) | 0.000002 | 1.5 |
| PPA | .013(.024) | 0.026 | 0.542 |
|  | Image-Word |  |  |
|  | Mean (Standard deviation) | *p*-value | Effect Size (*d*) |
| IPS | .19(.17) | 0.00003 | 1.12 |
| EVC | .015(.0035) | 0.061 | 0.43 |
| OPA | .068(.049) | 0.000005 | 1.39 |
| PPA | .011(.020) | 0.019 | 0.55 |

To visualize whether format decoding was similar across IPS and OPA, each ROI’s neural RDM was submitted to multidimensional scaling (MDS), which are projected into two-dimensional maps of each spatial direction and format. In these maps (Figure 6C), each arrow depicts one trial type, and the distance between arrows can be interpreted as the pair’s representational dissimilarity. For ease of interpretation, and to be consistent with the spatial decoding in IPS described in the spatial direction similarity analysis, we collapsed across left, right, and ahead. The MDS plots emphasize that while both regions distinguish between all three formats, schemas and words are more clearly disambiguated in IPS. Notably, format accounts for a large proportion of the variance captured by both regions, in spite of the fact that participants were asked to respond only to the spatial direction in the stimulus independent of the format.

#### Format decoding in searchlight analyses

Format decoding was robust within our regions of interest. We also queried the whole brain. We ran two searchlight analyses to see where formats were decoded across the whole brain. First, we analyzed which regions represented images as more similar to images than images to schemas or words. These regions are visualized in hot colors in Figure 7. In addition to parietal lobes, canonical scene regions (OPA, RSC, PPA) have higher correlations between images than with images to other formats. Second, we analyzed which regions represented schemas as more similar to schemas compared to words, and words more similar to words than schemas. This analysis uses the same baseline, word-schema correlations, and thus cannot distinguish whether these regions represent words as more similar to words, schemas more similar to schemas, or both. These regions are visualized in cool colors in Figure 7. Here, we saw bilateral fusiform gyrus, and inferior lateral occipital cortex, regions which have been implicated in word and object processing.

**Figure 7.**
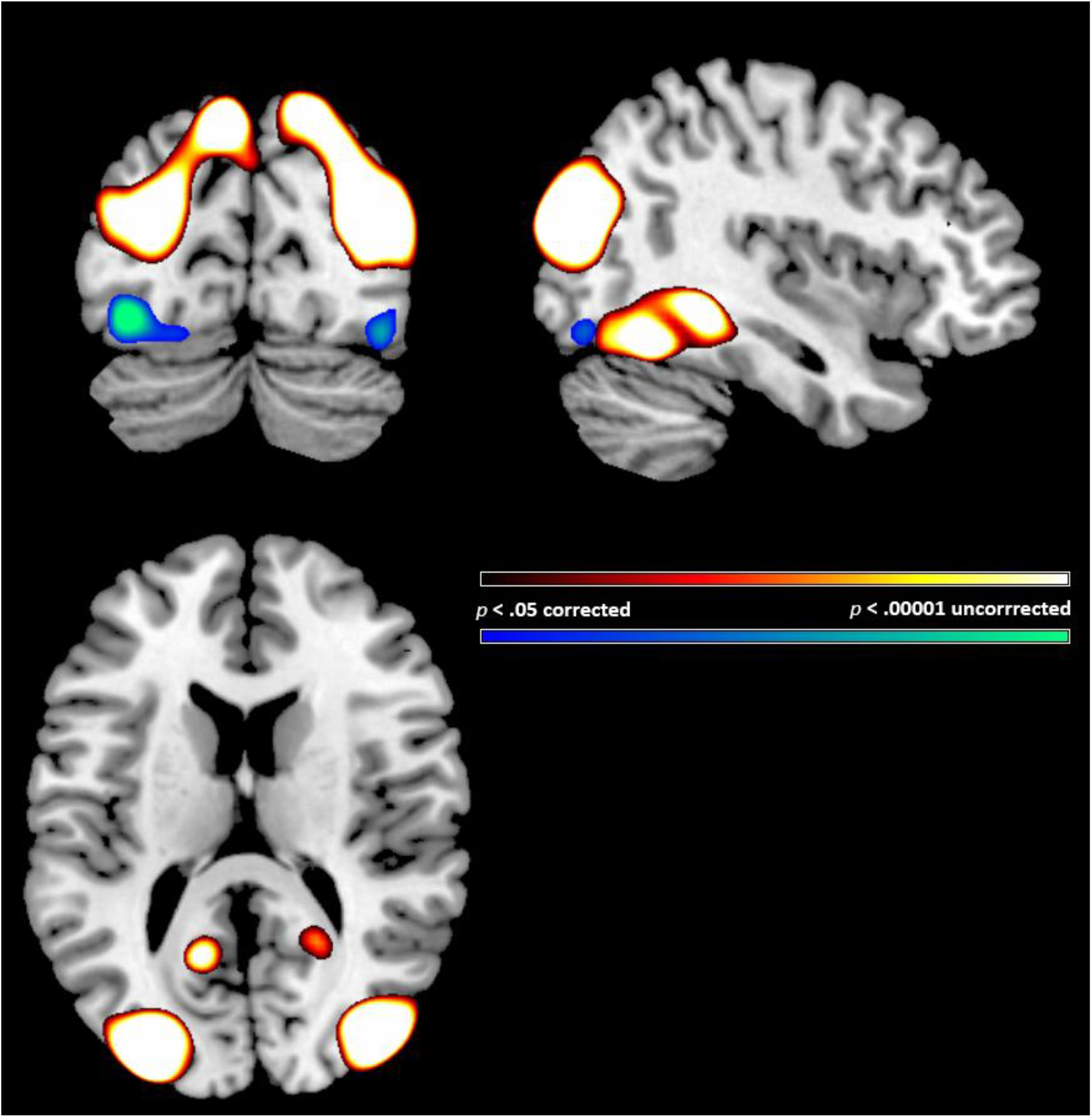
Decoding of format, whole brain searchlight. The theoretical RDM from Figure 5A generated the contrast between same format minus different format correlations for images-images minus images-words/schemas (in hot colors) and for schemas-schemas and words-words minus schemas-words (in cool colors). Image correlations were strongest in scene regions (OPA, PPA, RSC) and IPS, whereas schema and word correlations were strongest in word and object visual areas. Lower bound for searchlights are permutation corrected thresholds; upper bounds are *p* < .00001 uncorrected.

#### Individual differences in cognitive style

In an exploratory analysis, we correlated both dimensions of the Verbalizer-Visualizer Questionnaire (VVQ) with spatial direction decoding, within and across formats, in IPS. Correcting for multiple comparisons, no significant spatial direction decoding correlations were found. This question would be addressed more appropriately with a larger sample size. Responses to the de-briefing questionnaire also indicated that some participants preferred to say words to themselves, whereas others preferred to picture directions, or even imagine part of their body (e.g., their left shoulder for slight left). This variability in self-reported strategy suggests individual differences in cross-format decoding that a higher-powered study could address.

## Discussion

We aimed to investigate how spatial directions conveyed by distinct representational formats – visual scenes, schemas, and words – are behaviorally processed and neurally organized. We hypothesized that schemas and words elide spatial processing required by visual scenes and are processed more efficiently. This work bridges non-human models of navigation and cognitive mapping from visual scenes (Poucet, 1993; Etienne et al., 1996; Chen et al., 2013), with human research, which can investigate schematic and verbal communication of spatial directions.

Our findings support a model of spatial direction processing which taps a network that computes paths in visual scenes (Schindler & Bartels, 2013), but eschews in-depth spatial computations for efficient format-specific visual processing. Computing spatial directions from visual scenes requires imagining travel on paths shown. Computing spatial directions from words and schemas requires only visual identification. Visual scenes contain concrete detail, irrelevant to the spatial direction, but allow navigators to imagine traveling through the scene. By contrast, schemas and words contain easily distinguishable abstract direction information, but do not invite imagining travel in the same way as scenes.

In support of this model, we report three main findings. First, people responded to schemas and words more quickly than to scenes. Second, the intraparietal sulcus (IPS) bilaterally decoded spatial directions in scenes (and the decoding was structured into spatial categories, rather than arbitrary categories), and across the three formats, but not within schemas or words. These two findings suggest that, compared to words and schemas, scenes require relatively costly spatial computation to decode spatial directions in IPS. Third, format decoding independent of spatial directions was robust in ROI and whole-brain searchlight analyses. This finding suggests that, despite being task irrelevant (i.e., once the spatial direction is encoded, participants are better off discarding format, since spatial direction can be queried in any format), formats tap distinct neural pathways to convey relevant information.

Why might scenes be processed more slowly than schemas and words? First, unlike schemas, which discard irrelevant visual information and distill conceptually-important content, visual scenes contain detail unnecessary to compute the direction being depicted. Second, directions conveyed by schemas (at least in the current experiment) and words contain the exact same information about the spatial direction. Visual scenes can deviate from, for example, an exact 90° left turn. Thus, the direction in a visual scene must be computed for each presentation, then compared to the previous stimulus, whereas schemas and words need not be processed with this level of discrimination.

If spatial directions are computed from visual scenes, brain regions which support direction processing should contain representations of spatial directions for visual scenes, but not for schemas or words. This pattern was observed in the IPS bilaterally, regions of the brain implicated in egocentric spatial direction processing (Karnath, 1997; Whitlock et al., 2008; Galati et al., 2010; Schindler and Bartels, 2013).

We also found cross-decoding between schemas, words, and visual scenes in the IPS bilaterally. One explanation of our results is that when an individual views a scene, the IPS compute egocentric spatial directions from visual scenes by imagining the path of travel, resulting in a strong signal for each direction. However, when an individual views a schema or word, discerning spatial direction does not require IPS to compute egocentric spatial directions, yet it does so transiently, resulting in a weak signal. Within schema and word formats, this weak signal might not itself be decodable. Comparing the weak signal from schemas to the strong signal from scenes could yield cross-format decoding.

We did not observe spatial direction decoding in OPA, PPA, or RSC. The current results are not necessarily at odds with previous research showing spatial direction decoding in OPA (Julian et al., 2016; Bonner and Epstein, 2017) because our participants did not view walkable pathways. The OPA is causally involved in representing spatial directions defined by visual scene geometry and boundaries (e.g., constrained by hallways, counters, etc.) for obstacle avoidance. The lack of spatial direction decoding in PPA aligns with the hypothesis that the PPA codes a local visual scene (Epstein, 2008) in a viewpoint-invariant manner (Epstein et al., 2003). This hypothesis is supported by data showing that the neural representation in PPA is consistent across the same viewpoint of the same scene, but different when the viewpoint changes, suggesting that the PPA encodes spatial direction only with respect to the same scene. PPA may not have decoded directions in our experiment, because our examples used different scenes. Similarly, we can reconcile the current results with research showing allocentric spatial direction decoding in RSC (Vass and Epstein, 2013; Marchette et al., 2015). RSC contains representations of (allocentric) spatial directions that are aligned with respect to a prominent direction in the environment (e.g., the major axis of a building (Marchette et al., 2015); or the direction of a distal landmark, like a city in the distance (Shine et al., 2016)). IPS, on the other hand, contains representations of (egocentric) spatial directions that are aligned with respect to an individual’s current facing direction (Schindler & Bartels, 2013). Participants encoded directions in the current experiment egocentrically, supported by behavioral evidence – reaction time correlated with egocentric angle between the current and preceding spatial directions (but not visual angle). We do not know if changing task instructions to promote allocentric direction coding (e.g., asking participants to encode the spatial direction with respect to different sides of the screen), would yield cross-format spatial direction decoding in RSC.

Do schemas occupy a middle ground between words (abstract and arbitrarily related to the concept they denote), and visual scenes (concrete, rich in relevant and irrelevant detail)? Other conceptual domains support this notion of neural overlap between schemas, words, and visual depictions of concepts. Previous work on spatial prepositions report neural overlap in regions which process schemas and words, and separate areas which process schemas and visual images (Amorapanth et al., 2012). Viewing action words (like running) and schemas also resulted in cross-format decoding in action simulation and semantics areas (Quandt et al., 2017). In the current work, cross-format decoding of spatial directions was present in brain regions that process egocentric spatial directions.

Despite format being irrelevant for the task, format decoding was robust. Whereas images were processed distinctly from schemas and words in visual scene regions, schemas and words were disambiguated in IPS, as well as in object and visual word form areas. This pattern of results supports a model of concept coding in which abstract features are extracted from stimuli in format-dependent regions, then conveyed to brain regions which perform computations on the abstract concept. This finding is consistent with our behavioral data, suggesting implicit neural differences in the way scenes, schemas, and words are processed.

One limitation of our results is that we cannot account for all task-based effects (Harel et al., 2014) such as requiring that spatial directions be grouped into seven categories or that participants must respond to any of the three representational formats. We used a naturalistic task because of its applied relevance. When reading directions, for example, one might need to match a ‘slight left’ from memory to an egocentric road direction, a task which is comparable to our rolling one-back design, and requires a navigator to translate words to scenes. Still, spatial directions are not always categorized discretely. During walking a human navigator can easily turn 145° clockwise, while not necessarily categorizing this turn as “sharp right.” Nevertheless, we observed spatially-specific categorization in bilateral IPS for visual scenes: lefts were more similar to each other than rights, excluding the exact same direction. Note that such processing is counter-productive for the one-back task. Representing slight left as more similar to left than to slight right means it is harder to disambiguate a slight left from a left.

In sum, the current experiments reveal similarities and differences in formats of spatial direction depictions. Behaviorally, people responded to schemas and words more quickly than visual scenes. Neural decoding of spatial directions for visual scenes occurred in IPS bilaterally. This region revealed evidence of cross-format, abstract representation of spatial directions. These data challenge the specificity of IPS in encoding egocentric spatial directions, and support a model of spatial processing wherein images involve spatial direction computation, whereas schemas and words do not.

## Acknowledgements

The authors wish to acknowledge NIH grants F32DC015203 to S.M.W., and R01DC012511 and NSF Science of Learning Centers award 1041707 (subcontract 330161-18110-7341) to A.C. They also acknowledge Sam Trinh for stimuli development and pilot testing, Antonio Nicosia for behavioral study data collection, and Russell Epstein for helpful comments and feedback.

## Footnotes

1 We are agnostic about whether the images used in this study can actually be considered visual scenes. We use the term “image” below to be consistent with the general terms ‘word’ and ‘schema.’ In the introduction and discussion, on the other hand, we discuss ‘visual scenes’ to connect the domain specific work here with other research on visual scenes. The robust activation of the scene network while participants viewed the images also lead us to speculate that participants treated these stimuli as scenes.

